# Amphetamine in Adolescence Induces a Sex-Specific Mesolimbic Dopamine Phenotype in the Adult Prefrontal Cortex

**DOI:** 10.1101/2025.02.26.640363

**Authors:** G. Hernandez, J. Zhao, Z. Niu, D. MacGowan, T. Capolicchio, A. Song, S. Gul, A. Moiz, I. Herrera, J. J. Day, C. Flores

## Abstract

Drugs of abuse in adolescence impact brain maturation and increase psychiatric risk, with differences in sensitivity between males and females. Amphetamine in adolescent male, but not female mice, causes dopamine axons intended to innervate the nucleus accumbens and to grow ectopically to the prefrontal cortex (PFC). This is mediated by drug-induced downregulation of the Netrin-1 receptor DCC. How off-target dopamine axons function in the adult PFC remains to be determined. Here we report that males and females show place preference for amphetamine in adolescence. However, only in males, amphetamine increases PFC dopamine transporter expression in adulthood: leading to aberrant baseline dopamine transients, faster dopamine release, and exaggerated responses to acute methylphenidate. Upregulation of DCC in adolescence, using CRISPRa, prevents all these changes. Mesolimbic dopamine axons rerouted to the PFC in adolescence retain anatomical and functional phenotypes of their intended target, rendering males enduringly vulnerable to the harmful effects of drugs of abuse.

## Introduction

Adolescence is a period of extensive biological and behavioral transformation, including profound reorganization of dopaminergic connectivity within the prefrontal cortex (PFC)^1,2^. As a hub for executive function, the PFC matures slowly, showing changes in structure and function well into adulthood^3–6^. The protracted development of the PFC is accompanied by an increase in the density of dopamine innervation^7–10^ which results from dopamine axons still extending from the nucleus accumbens (Nac) to the PFC^11–13^. This delayed growth is the only documented instance of long-distance axonal pathfinding in adolescence^12,14^ and depends on the precise signaling of the guidance cue receptor DCC (Deleted in Colorectal Cancer) and its ligand Netrin-1^12,15^.

Exposure to recreational-, but not therapeutic-like doses of AMPH in early adolescence decreases the expression of *Dcc* mRNA^15^ and DCC protein^16^ in dopamine neurons as well as of Netrin-1 protein levels in the Nac^15,17^. These disruptions cause mistargeting of dopamine axons, leading to the rerouting of mesolimbic projections to the PFC and to a reduced number of dopamine axon varicosities in this region^12,15,18^. These changes trigger alterations in the dendritic architecture of PFC pyramidal neurons, resulting in enduring impairment in inhibitory control^15,19^. Notably, AMPH-induced disruption of adolescent dopamine and cognitive development is sexually dimorphic – it occurs in male mice only^15^. Furthermore, CRISPRa-mediated upregulation of DCC receptors in the ventral tegmental area (VTA) in early adolescence in male mice induces female-like protection against AMPH-induced axon mistargeting and cognitive control dysfunction^15^.

Mesolimbic and mesocortical dopamine projections have distinct anatomical and functional characteristics^20–22^. Here we assessed if AMPH-induced disruption of dopamine axon pathfinding in adolescence impacts PFC dopamine dynamics in adulthood and if ectopic mesolimbic dopamine axons retain their anatomical and functional phenotype or adopt properties of their off target ^20,23,24^. We addressed these questions by combining behavioral and quantitative neuroanatomical analyses, dopamine dynamics monitoring, and CRISPRa gene editing in male and female mice.

## Methods

### Animals

Experimental procedures were performed according to the guidelines of the Canadian Council of Animal Care and approved by the McGill University/Douglas Mental Health University Institute Animal Care Committee. C57BL/6J mice were obtained from Charles River Laboratories (Saint-Constant, QC, Canada). All mice were maintained on a 12-h light–dark cycle (light on at 0800 h) in a temperature controlled (21C) facility with 42% humidity and given ad libitum access to food and water. Male and female mice were housed with same-sex littermates throughout the experimental procedures.

### Drugs and dose

d-Amphetamine sulfate (Sigma-Aldrich, Dorset, United Kingdom) was dissolved in 0.9% saline. All AMPH injections were administered i.p. at a volume of 0.1 mL/10 g. AMPH at a recreational-like dose (4 mg/Kg) was used across all the experiments^15,25^.

Methylphenidate (Apotex, Canada) was dissolved in 0.9% saline, and was administered i.p. at a volume of 0.1 mL/10 g. A single dose of 10 mg/kg was selected because it reliably increases extracellular dopamine levels in the PFC^26–28^.

### Stereotaxic surgeries

#### Fibre Photometry

Adult wild-type male and female (PND81+/−1) mice were anesthetized with Isoflorane (Fresenius Kabi, Canada) and then placed in a stereotaxic frame for microinfusion of the viral construct using a Stoelting Quintessential Stereotaxic Injector attached to a Hamilton 5µL microsyringe 75RN. An Adeno-associated virus (AVVs) expressing hSyn-rDA2h (Grab_DA2h_)^29^ (Canadian Neurophotonics Platform Viral Vector Core Facility RRID:SCR_016477) was injected (0.5 µL, tier 4×10^−12^ vg/mL) unilaterally into the left infralimbic region of the PFC using the following coordinates: AP: +1.8 mm relative to Bregma, ML: 0.3 mm DV - 2.5 from the skull. The infusion was done over eight minutes. Ten minutes after the end of the microinfusion, the needle was removed and an optical fiber (Doric Lenses, diameter 4.8; NA 0.66 Cat # MFC 400/430) within a zirconia ferrule was implanted into the PFC with its tip targeting 0.1mm above the injection coordinates. Ferrules were secured to the skull using dental cement with the help of two skull-penetrating screws. Mice were left undisturbed in their home cages for 3-weeks to maximize viral expression ^29,30^. Mice were group-housed during recovery and across the experiments.

### CRISPRa

#### (Clustered Regularly Interspaced Short Palindro mic repeats activation)

To increase DCC receptor expression in adolescence, we employed CRISPRa with 4 compatible single guide RNAs (sgRNAs) as in ^15^. Robust neuronal expression was ensured using lentiviral constructs optimized for this purpose (lenti SYN-FLAG-dCas9-VPR, RRID:Addgene_114196; lenti U6-sgRNA/EF1a-mCherry, RRID:Addgene_114199)^31^. A non-targeting sgRNA against the bacterial *LacZ* gene served as a control. For detailed information about construction and validation see ^15^.

Early adolescent wild-type C57BL/6 (PND21+/−1) male mice were anesthetized with Isoflorane (Fresenius Kabi, Canada) and then placed in a stereotaxic frame to bilaterally microinfused a total of 1.0 μl of lentiviral mix containing 0.33 μl of four *Dcc* sgRNAs or 0.33 μl of *LacZ* sgRNA and 0.66 μl of the dCas9-VPR virus in sterile PBS^15,31^. Using the Stoelting Quintessential Stereotaxic Injector attached to a Hamilton 5µL microsyringe 75RN. The lentiviral mixture was microinjected into the ventral tegmental area (VTA) using the following coordinates: AP: −2.56 mm relative to Bregma, ML: 0.75 mm DV - 4.5 from the skull at 10-degree angle. The infusion was done over 16 min. 10 minutes after the end of the infusion the needle was removed. Animals were left to recover for 24 hours before starting the place preference conditioning.

### Conditioned Place preference

Male and female mice were separately assessed in a biased conditioned place preference (CPP) paradigm using 4 mg/kg AMPH or saline during early adolescence (PND 21-32), as previously described^15,32^. On PND20, each mouse freely explored the CPP apparatus (two distinct chambers connected by a neutral area) for 20 minutes. Initial chamber preference was determined by calculating the percentage of time spent in each chamber. The less preferred chamber was then paired with AMPH (experimental group) or saline (control group) in alternating 30-minute conditioning sessions over 10 days (5 AMPH, 5 saline). Control mice received saline exclusively. On PND32 and PND80, a 20-minute conditioning test was conducted to assess chamber preference. Delta preference score (Δ Place Preference) was calculated as: % time spent in the initially unpreferred chamber (post-test) – % time spent in the same chamber (pre-test).

### Fiber Photometry

Fiber photometry signals were collected using a fiber photometry rig with optical components from Doric. The LED beams were reflected and coupled to a fluorescence minicube (FMC4, Doric Lenses). A 0.5 m-long optical fiber (400mm, Doric Lenses) was used to transmit light between the fluorescence minicube and the implanted fiber. Lenses were controlled by a real-time processor from Tucker Davis Technologies (TDT; RZ10). TDT Synapse software was used for data acquisition. 465 nm and 405 nm LEDs were modulated at 211Hz or 230 Hz and at 330 Hz, respectively. LED currents were adjusted to return a voltage between 150 and 200 mV for each signal, were offset by 5 mA, and were demodulated using a 4 Hz low-pass frequency filter. During the experiment, mice were placed in an open field (11-1/2″ Long x 7-1/2″ Wide x 5″ Deep) and attached to the optical fiber. First, they were allowed to habituate to the environment for 1h. Following a 10-minute baseline recording, mice received an intraperitoneal (i.p.) injection of saline, and recording continued for 20 minutes. Mice were then administered an i.p. injection of methylphenidate (10 mg/kg), and recordings were continued for an additional 90 minutes.

### Neuroanatomical Analysis

#### Perfusion

In adulthood and at the end of the experiment (PND101 ± 15), mice were anesthetized with a cocktail of ketamine 100 mg/kg, xylazine 10 mg/kg, acepromazine 3 mg/kg and perfused with cold PBS followed with 4% paraformaldehyde. The brains were sliced into 35μm coronal sections using a Leica vibratome.

#### Immunolabeling

As previously done, every second coronal brain section was processed for immunofluorescent labeling (1:2 series)^33^. Tissue sections were blocked (2% bovine serum albumin, 0.2% Tween-20 in PBS) for 1 hour and subsequently incubated for 48 hours at 4 °C with a monoclonal rat anti-dopamine transporter (DAT) antibody (1:500, MAB369, Millipore Bioscience Research Reagents) and a polyclonal rabbit anti-tyrosine hydroxylase (TH) antibody (1:500, AB152, Millipore Bioscience Research Reagents) diluted in blocking solution. Sections were then rinsed with 0.2% PBS-T, and then incubated for 2 h at room temperature with Goat anti-Rat Alexa Fluor 488 and Donkey anti-Rabbit Alexa Fluor 594-conjugated secondary antibodies (1:500; Invitrogen, A-11006 and A-21207). Sections were coverslipped onto gelatin-coated slides using a SlowFade Gold Antifade mounting medium (Invitrogen).

#### Stereology

The cingulate 1 (Cg1), prelimbic (PrL), and infralimbic (IL) subregions of the pregenual PFC were delineated according to plates spanning 14–18 of the mouse brain atlas (Paxinos and Franklin, 2019) and contours were traced at 5X magnification using a Leica DM400B microscope along the dense TH-positive innervation of PFC layers V-VI ^34^. We assessed the density of DAT+ varicosities in the Cg1, PrL and IL subregions using a stereological fractionator sampling design ^35^ Stereoinvestigator software (MBF, St. Albans VT) as previously ^12,34^. Varicosities were counted at 100X magnification.

#### Verification of viral transduction CRISPRa

Every second section was processed for visualization of tyrosine hydroxylase (TH)+ neurons and mCherry – the reporter tag added to the sgRNA. Sections were rinsed 3 times for 10 min with 1x PBS and blocked in 2% bovine serum albumin (in 1x PBS and Tween-20) for 1 h at room temperature. Sections were then incubated for 48 h at 4 °C in the primary antibodies: mouse anti-TH (Millipore Sigma, cat. no. MAB318) and rabbit anti-red fluorescent protein (RFP) for mCherry (Rockland, cat. no. 600-401-379). Sections were rinsed 3 times for 10 min with 1x PBS and incubated in the secondary goat anti-mouse Alexa Fluor 488 antibody (Invitrogen, cat. no. A-11001) and the secondary donkey anti-rabbit Alexa Fluor 594 antibody (Invitrogen, cat. no. A-21207), for 1 h at room temperature. Sections were rinsed 3 times in 1x PBS and mounted with VECTASHIELD Hardset antifade mounting medium with DAPI (Vector Laboratories, cat. no. H-1500-10). Representative images were taken using an epifluorescent microscope (Leica DM400X3).

To label GRAB_DA_, tissue sections were rinsed and then immunostained with chicken anti-GFP antibody (1:500, Abcam, Cat#ab13970), followed by the Alexa-488-conjugated goat-anti-chicken (1:200, AAT-Bio, Cat#16687) secondary antibody.

### Data analysis

*CPP data* were analyzed using a three-way repeated measures ANOVA with conditioning test Day (PND32; PND80) as the repeated measure and drug treatment (AMPH; saline) and sex as between subjects factors. Due to the difference in variance in the CPP data obtained in the CRISPRa experiment, these data were analyzed using Generalized Estimating Equations (GEE) with two-independent (sgRNA and Treatment) and one repeated (conditioning tests) factors.

*Density of DAT+ varicosities data* were analyzed using a three-way repeated measures ANOVA, with PFC subregion as the repeated measure and drug and sex as between subjects factors.

*Dopamine dynamics data* were processed using a MATLAB script (Supplementary Materials). Noise-related changes in *GRAB*_*DA*_ fluorescence across the whole experimental session were removed by scaling the isosbestic control signal (405nm) and regressing it onto the dopamine-sensitive signal (465nm). This regression generated a predicted model of the noise that was based on the isosbestic control^36^. Dopamine-independent waveforms on the 405nm model were then subtracted from the raw Grab_DA2h_ signal. The resulting signal was converted to ΔF/F0 by dividing it to the fitted isosbestic control.

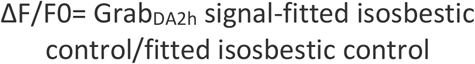

A linear regression to correct for photobleaching was fitted to data obtained before vehicle injection and applied across the series. For comparison of data across mice, a robust z-score^37^ (z ΔF/F0) was computed using the median and the median absolute deviation (MAD) from the 10 minutes preceding the saline injection.

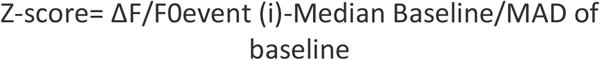

We calculated the area under the curve and used a two-way ANOVA to quantify the effect of treatment (saline or AMPH) and sex.

The baseline transient frequency and amplitude were processed using a MATLAB script (supplementary material) and the peak finder procedure written by Nathanael Yoder (2023). This function searches noisy signals for derivative crossings (local maxima/minimal) that are at least a specified amount above or below the last derivative crossing. Peak amplitudes were measured from the last local minimal found by the function.

The dopamine signal observed after the i.p. saline injection was measured using exponential equations that were fitted to the ascending and descending parts of the curve using MATLAB. The ascending part of the curve was fitted to the following equation:

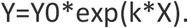

The descending part was fitted to the one phase decay equation:

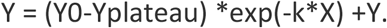

In these equations Y0 is the Y value when X (time) is zero; Yplateau is the value of Y at infinite times, k is the rate constant.

The parameters obtained were then analyzed using a two-way ANOVA with treatment in adolescence (saline, AMPH) and sex as independent factors.

In all the statistical analyses, Holm-Šídák’s multiple comparisons tests were used when significant interactions were detected.

All statistical tests are described in Tables 1–5. All statistical analyses were carried out using Prism software (GraphPad), except for the GEE analysis, which was done in SPSS.

## Data Availability

The data supporting the finding as the MATLAB code are openly available in g node doi: https://doi.org/10.12751/g-node.qq93iz and figshare https://doi.org/10.6084/m9.figshare.28489385

## Results

### AMPH in adolescence increases the density of DAT+ axons in the adult PFC of male, but not female mice

Mesolimbic and mesocortical dopamine axons have distinct anatomical, molecular and functional properties^20,21^. In male mice, early adolescent exposure to AMPH reroutes Nac dopamine axons to the PFC^15^. To test whether ectopic mesolimbic dopamine axons retain the molecular phenotype of their intended target or adopt those of the PFC, we quantified the density of DAT+ varicosities in the adult PFC of mice exposed to AMPH or saline in early adolescence (Fig. 1 A). We performed this experiment in male and female mice to assess for potential sex differences.

**Figure 1.**
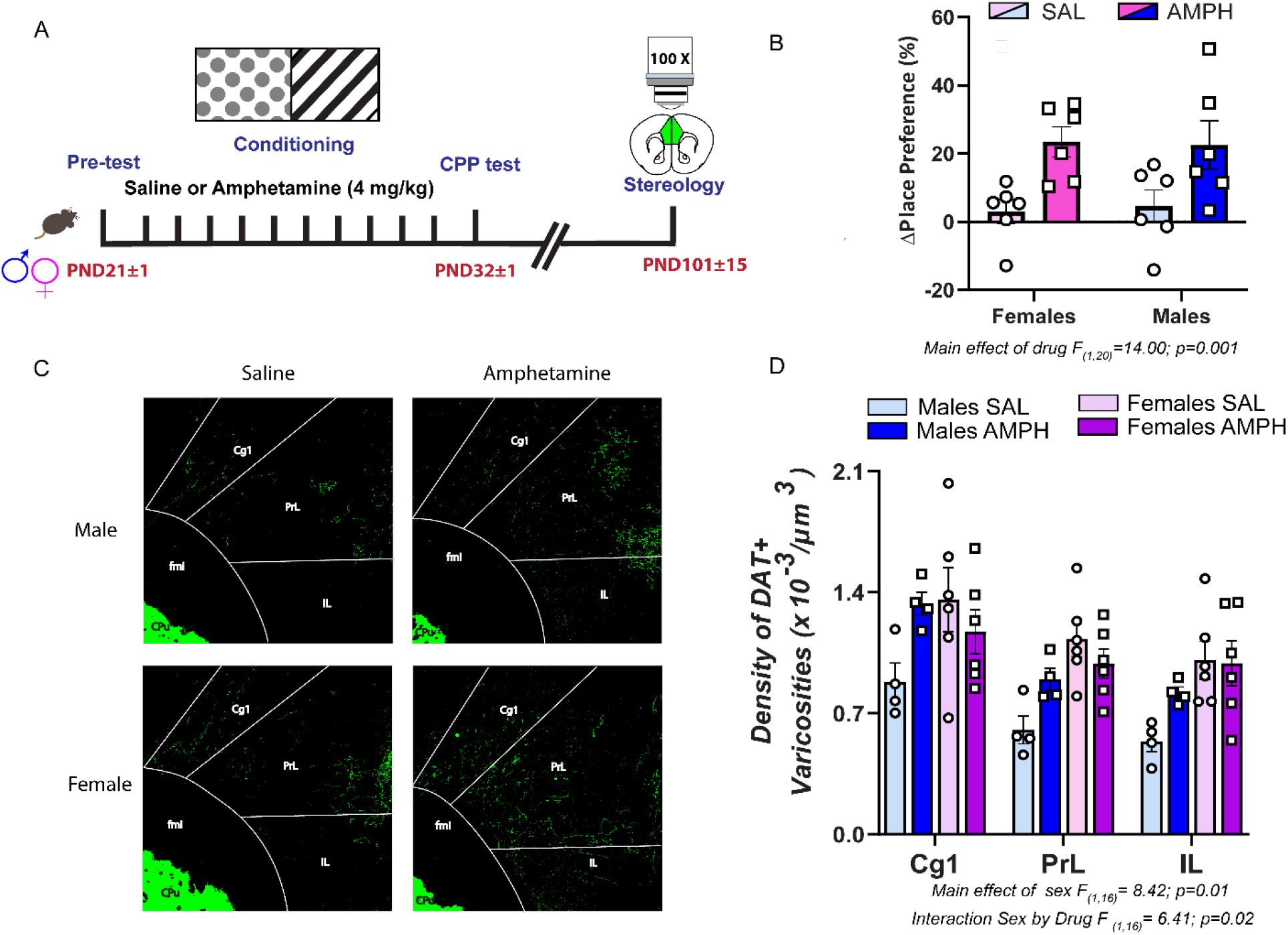
Rewarding doses of amphetamine (AMPH) in adolescence increase the density of dopamine transporter (DAT)+ axons in the adult PFC of male, but not female mice. **A**. Experimental timeline. **B**. Male and female mice exposed to AMPH (4.0 mg/kg) in early adolescence develop place preference for the side of the box paired with the drug **C**. Photoimages of representative coronal section of DAT+ axons in the cingulate (Cg1), prelimbic (PrL) and Infralimbic (IL) subregions of the PFC of adult male and female mice exposed to AMPH or to saline in adolescence. In males, AMPH in adolescence increases the number of DAT+ axons compared to saline controls (*Top)*. Adult females display similar levels of DAT+ axons regardless of adolescent treatment (*Bottom)*. In saline control groups, females exhibit higher number of DAT+ varicosities than males across the three PFC subregions. **D**. Stereological analysis revealed increased density of DAT+ varicosities adult males exposed to AMPH in adolescence compared to their saline counterparts. In females, no differences between AMPH and saline groups were found. In saline pre-treated groups, adult females have higher density of DAT+ varicosities, compared to males. All bar graphs are presented as mean values ±SEM. Source data are provided as a Source Data file.

AMPH in early adolescence induces robust and comparable levels of CPP in both males and females, 24 h after the last conditioning session, in line with our previous report15 (Fig. 1B). In adulthood, AMPH-pretreated males but not females show a robust increase (∼50%) in the density of DAT+ varicosities across the Cg1, PrL and IL PFC subregions, compared to their saline counterparts (Fig 1 C&D). This indicates that misrouted mesolimbic dopamine axons maintain molecular signatures of their intended target.

Notably, in the saline control groups, females show higher PFC DAT+ varicosities density (55%) compared to males. To our knowledge, this is the first demonstration of increased DAT expression in the PFC of males compared to females. Our findings suggest sex differences in dopamine reuptake in this region, which may translate into differences in PFC function and cognitive processing.

### AMPH-induced CPP incubates during the adolescent period

We next examined if AMPH in early adolescence leads to long-term CPP and if this effect is sex-specific (Fig 2A). As expected, AMPH treatment induced preference for the drug-paired compartment 24 hours after the last conditioning session. This preference persisted and became more prominent in adulthood (Fig. 2B), revealing an incubation-type effect and suggesting that the rewarding effects of AMPH are latent and strengthen with age^38^.

**Figure 2.**
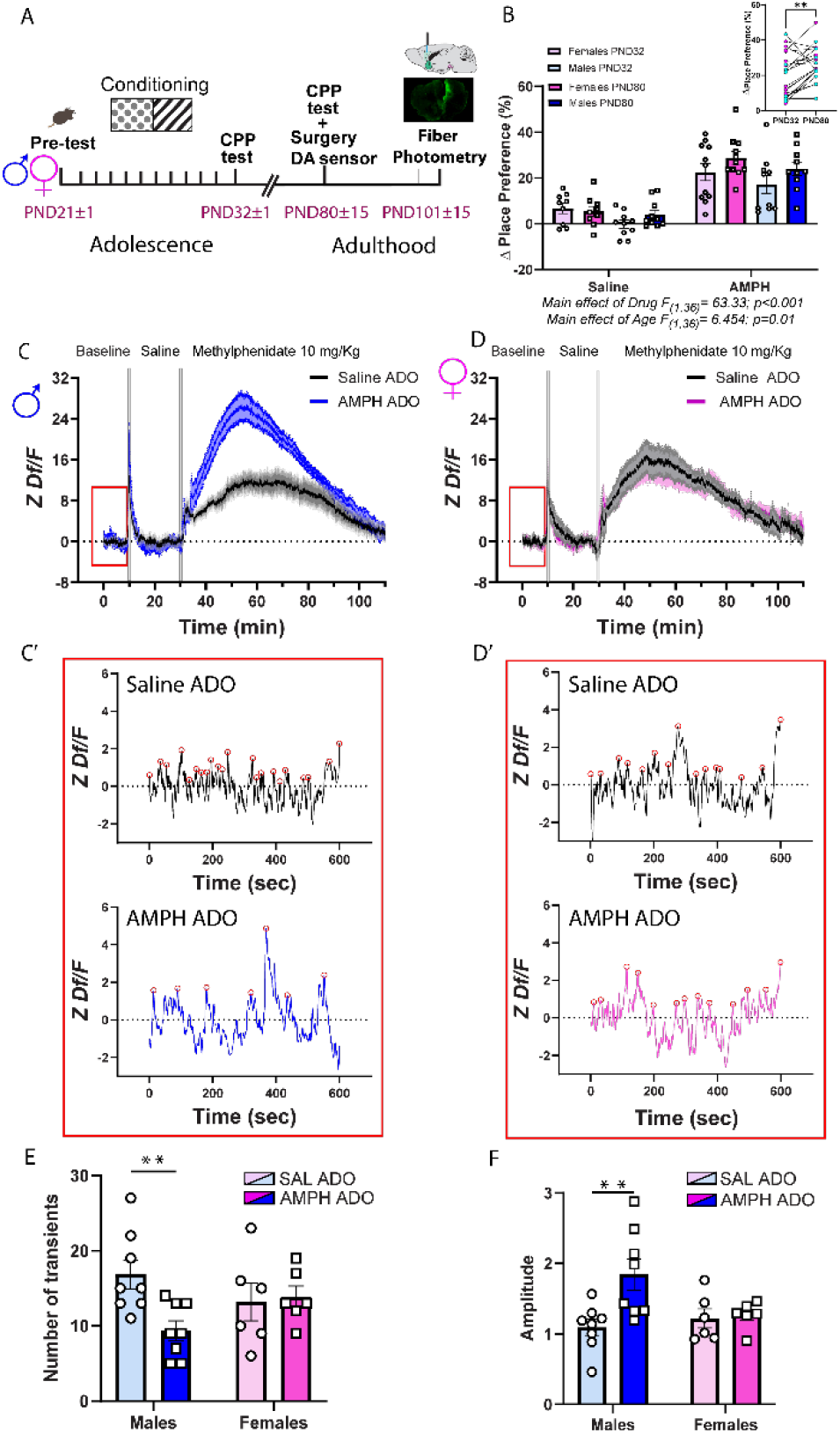
In males only, AMPH in early adolescence increased PFC dopamine accumulation at baseline in adulthood. **A**. Experimental timeline. **B**. Male and female mice exposed to AMPH (4.0 mg/kg) in early adolescence developed place preference for the side of the box paired with the drug. This effect increased over time, suggesting an incubation-like effect (inset). **C**. Average PFC dopamine dynamics during baseline and following an acute i.p. injection of saline and of methylphenidate. The red rectangle highlights the baseline period used in subsequent analyses. **C’-D’**. Representative dopamine traces from adult males and females exposed to saline or AMPH in adolescence. Red circles indicate individual dopamine transients **E**. Males administered AMPH in adolescence show reduced number of dopamine transients during baseline. This reduction is not seen in females. **I**. The amplitude of dopamine transients is significantly greater in AMPH-treated males compared to their saline counterparts. Females displayed similar transient amplitude regardless of adolescent treatment. All bar graphs are presented as mean values ±SEM. Source data are provided as a Source Data file. *p<0.05; ** p<0.01. ADO = adolescence.

### AMPH in adolescence leads to exaggerated dopamine responsiveness in the adult PFC of males

To explore the functional consequences of altered PFC dopamine innervation in adult males exposed to AMPH in early adolescence, we investigated dopamine transient kinetics and dynamics using fibre photometry. Twenty-four hours after the adult CPP test, AMPH- and saline-pretreated male and female mice underwent stereotaxic unilateral microinfusion of AAV-hSyn-rDA2h (Grab_DA2h_) into the IL part of the PFC, followed by intracranial implantation of an optical fibre. Three weeks later, we quantified dopamine signals at baseline, following an i.p. injection of saline and following an acute i.p. administration of methylphenidate at a 10 mg/kg dose (Fig 2A).

#### Baseline

Using the results obtained from the peak finder procedure^39^ we compared the number and amplitude of dopamine transients during the ten-minute baseline, right after the 1h habitation period (Fig. 2C&D, red square). We found that adult male mice treated with AMPH in early adolescence have a reduced number of spontaneous dopamine transients compared to saline controls, most likely due to increased dopamine reuptake. This effect of AMPH is not observed in females. There are no sex differences in the baseline number of dopamine transients (Fig. 2E). Interestingly, in males, AMPH administered in adolescence *increases* baseline dopamine transient amplitude in adulthood (Fig. 2F). This may be the result of a combined effect of reduced transient frequency and increased amount of readily releasable dopamine due to higher density of DAT+ varicosities in the PFC. In females, there are no differences between AMPH and saline groups, and, surprisingly, there are no differences in baseline dopamine transient amplitude between males and females despite the sex differences in DAT expression (Fig. 2F).

#### Saline

If in AMPH-pretreated males, dopamine accumulation at terminals is enhanced during baseline conditions due to an increase in dopamine recycling, this group should show accelerated dopamine release and clearance in response to an acute i.p. injection of saline (Fig. 3A&B). We assessed this possibility by fitting a double logarithmic function to the ascending and descending parts of the fluorescent peak. Indeed, in adult males, an acute saline injection induces faster dopamine release in the AMPH-versus the saline-pretreated groups. Accelerated dopamine release is indicated by both the steeper slope (k ascending) (Fig. 3C) and the shorter time it takes for the fluorescence signal to double (doubling time) (Fig. 3D). In females, the timing of dopamine release does not differ between AMPH- and saline-pretreated groups.

**Figure 3.**
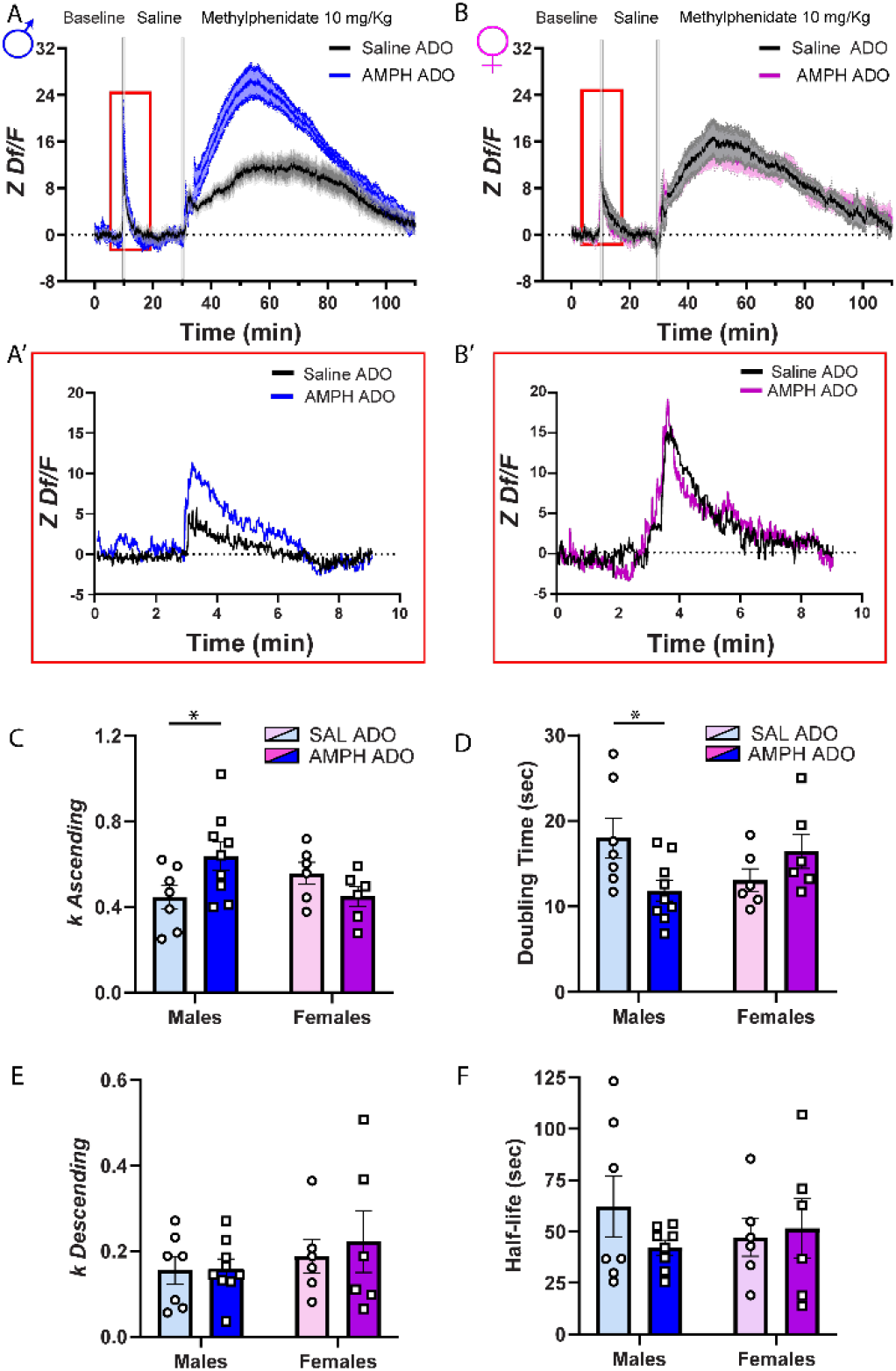
AMPH in adolescence accelerates PFC dopamine dynamics following a saline injection in adulthood, but only in males. **A**. Average PFC dopamine dynamics during baseline and following an acute i.p. injection of saline and of methylphenidate. The red rectangle highlights dopamine release following the saline challenge. **A’-B’**. Representative dopamine traces from adult males and females exposed to saline or AMPH in adolescence. Peak analysis was performed on each trace after the acute i.p. injection of saline. To each subject, we fitted two exponential curves one for ascending part and one for the descending part to calculate the slope of each peak (*k*); the time for the signal to double in size (doubling time) and the time the signal decreases in intensity by half (half-life). **C**. The *k* value for the ascending curve of AMPH-pretreated males is significantly higher than that of their saline counterparts, suggesting a faster surge of dopamine release in adult males exposed to AMPH in adolescence. In females, the slopes for the ascending curve are similar between treatment groups. **D**. Due to the more rapid increase in dopamine release in AMPH-pretreated males, the resulting doubling time is significantly shorter compared to that observed in males exposed to saline in adolescence. In females, the doubling time is similar regardless of adolescent treatment. **E**. The slopes observed in the descending curves of both male and female groups demonstrate a consistent pattern, indicating a comparable rate of decline in dopamine concentration across all groups. **F**. For the half-lives, there are no differences across groups. All bar graphs are presented as mean values ±SEM. Source data are provided as a Source Data file. *p<0.05. ADO = adolescence.

In both male and female mice, we were unable to capture any difference in the fluorescent signaling used to infer reuptake (FIG 3E-F) since both groups show similar slope (k descending) and half-life parameters. This result is surprising because we expected a faster dopamine clearance in AMPH-treated males due to their increased DAT expression. The high-affinity DA sensor used, which provides high affinity but has a slow signal decay time (8.3 sec) ^40,41^, could be masking the detection of rapid clearance events.

#### Methylphenidate

Previous work has shown that DAT overexpression increases methylphenidate-induced increase in extracellular dopamine, at least in the striatum^42^. One key role of DAT is to facilitate the reuptake of dopamine from the extracellular space back into the presynaptic terminal, thereby enabling its reuse^43^. However, dopamine terminals in the PFC are relatively inefficient at reuptake due to low DAT density^44^; with the norepinephrine transporter primarily handling this function^45,46^. This inefficiency results in reduced dopamine recycling and reserve pools, with most of the dopamine available for release being newly synthesized^47^.

We assessed dopamine signaling in male and female mice after a methylphenidate injection. We anticipated that DAT overexpression in males exposed to AMPH in adolescence would result in exaggerated methylphenidate-induced dopamine signaling due to the increased density of DAT+ varicosities in the PFC. We also anticipated sex-specific methylphenidate-induced dopamine signals considering the increased density in DAT+ varicosities in females compared to males.

Acute methylphenidate increased dopaminergic signaling across all groups. However, males that received AMPH in adolescence showed significantly higher dopamine release 10 min after the methylphenidate challenge. This signal peaked 30 min after the injection and returned to baseline levels 80 min after (Fig.4A). In contrast, female mice exposed to AMPH or saline in adolescence showed similarly elevated dopamine levels in response to methylphenidate (Fig. 4B). Analysis of the area under the curve confirmed that males pretreated with AMPH, but not females have exaggerated dopamine increase in response to methylphenidate (Fig. 4C). These findings indicate that ectopic dopamine innervation renders the male PFC hypersensitive to methylphenidate-induced dopamine release. Contrary to our prediction, there were no differences in dopamine signaling between saline-treated male and female groups.

**Figure 4.**
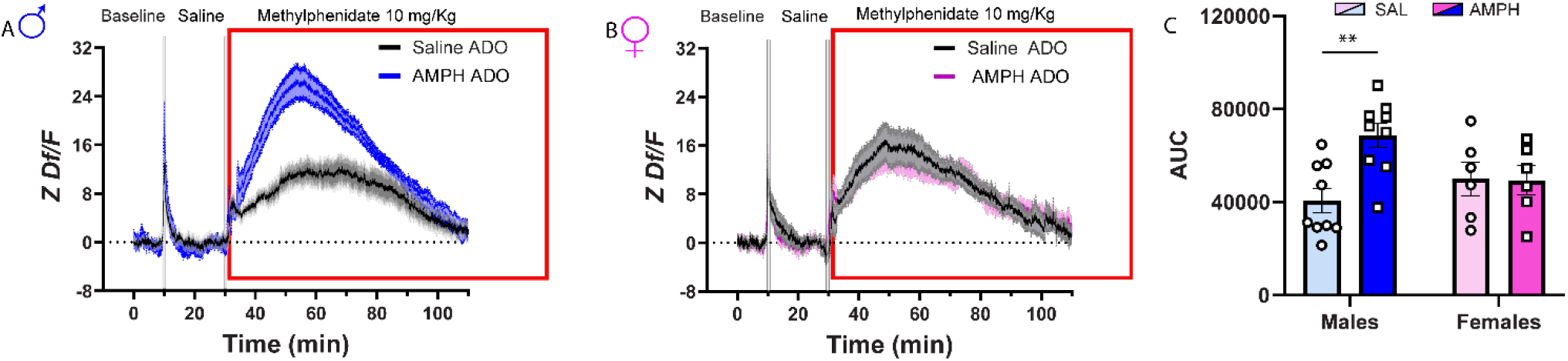
AMPH in adolescence leads to exaggerated methylphenidate-induced increase in PFC dopamine release in adult males only. **A**. Average PFC dopamine dynamics during baseline and following an acute i.p. injection of saline and of methylphenidate. The red rectangle highlights dopamine release following acute i.p. methylphenidate challenge. Methylphenidate (10 mg/Kg) increases PFC dopamine release in both sexes, but this effect is significantly amplified in males with early adolescent AMPH exposure **B**. In females, methylphenidate also increases PFC dopamine release, but this increase is similar in AMPH and saline groups. **C**. These observations are corroborated by quantifying the are under the curve (AUC). All bar graphs are presented as mean values ±SEM. Source data are provided as a Source Data file. **p<0.01. ADO = adolescence.

### DCC receptor upregulation in adolescence prevents AMPH-induced place preference in adolescence

We next evaluated whether increasing DCC receptor expression in adolescence, via CRISPRa, prevents changes in PFC dopamine dynamics observed in adult males exposed to AMPH in adolescence. As previously shown^15^, we used a combination of four *Dcc* sgRNAs which increases *Dcc* mRNA up to 4-fold in dopamine cell culture and produces a significant in-vivo increase in DCC protein expression in axons of dopamine neurons in the Nac, compared to *LacZ* sgRNA control infection^15^.

Male mice were injected with the sgRNA cocktail and dCas9 viruses at PND21 and then exposed to CPP. Place preference was assessed 24 h and 50 days after the last injection (Fig. 5A). After the last preference test, mice were microinjected with a virus expressing GrabDA2h and an optical fiber aimed at the IL area of the PFC. Fiber photometry assessment of dopamine release was done 3 weeks after.

**Figure 5.**
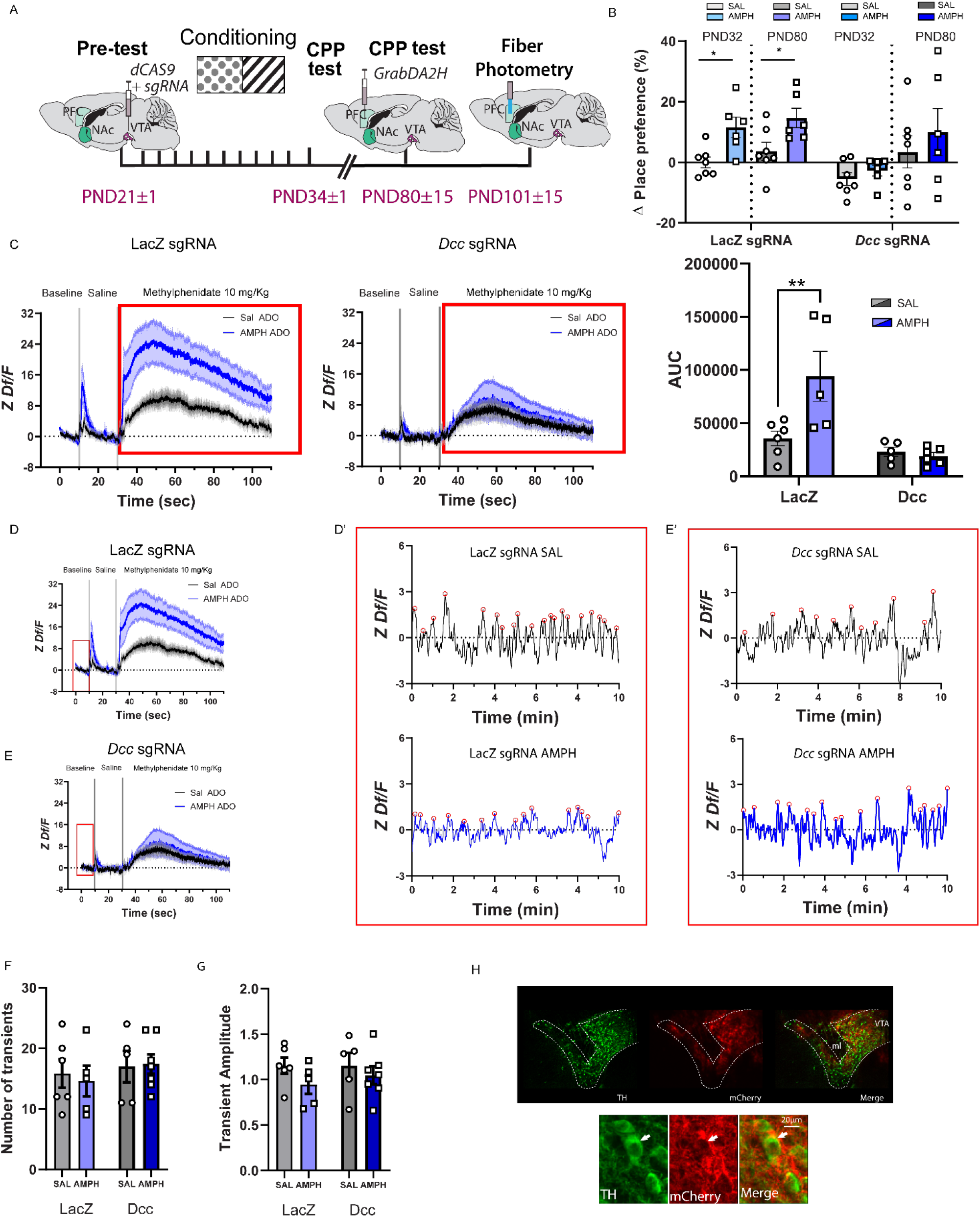
Increasing *Dcc* transcription in the VTA with CRISPRa prevents PFC dopamine dysfunction in adult males exposed to adolescent AMPH. **A**. Experimental timeline. **B**. AMPH induces conditioned place preference in the *LacZ* sgRNA group in adolescence and this preference persists into adulthood. Increased VTA *Dcc* transcription in adolescence impairs AMPH-induced place preference in adolescence and in adulthood. **C**. AMPH in adolescence increased adult PFC dopamine release in response to the methylphenidate (10 mg/kg) challenge in adulthood in the *LacZ* sgRNA group compared to their control counterparts. This effect was completely prevented in mice that had *Dcc* upregulation in adolescence. The area under the curve analysis corroborates this finding **D-E**. Average PFC dopamine dynamics during baseline and following an acute i.p. injection of saline and of methylphenidate in males with CRISPRa-induced *LacZ* (C) or *Dcc* (D) sgRNA transcription. The red rectangle highlights the baseline period used in subsequent analyses. **D’-E’**. Representative dopamine traces from *LacZ* and *Dcc* sgRNA groups exposed to AMPH or saline in adolescence. Red circles indicate individual dopamine transients. **F**. There is no difference in the number of transients during the baseline period between *LacZ* and *Dcc* sgRNA groups regardless of treatment in adolescence. **G**. There is no difference in the transient amplitude between *LacZ* and *Dcc* sgRNA-groups regardless of treatment in adolescence. **H**. Photoimages of coronal VTA sections showing low and high magnification of dopamine neurons (TH+ green) co-expressing the sgRNA viruses (mCherry). Arrowhead denotes colocalization. All bar graphs are presented as mean values ±SEM. Source data are provided as a Source Data file. **p<0.01. ADO = adolescence.

Adult male mice exposed to AMPH in adolescence and infected with *LacZ* sgRNA showed robust CPP 24h after the last conditioning session. The preference for the drug-paired compartment is maintained when mice reached adulthood. Remarkably, DCC receptor overexpression abolished AMPH-induced place preference both 24 h after the last conditioning session and when re-tested in adulthood, although higher preference variability emerged at PND80 (Fig. 5B). These results indicate a role of DCC receptors in AMPH-induced place preference and may be related to changes in drug-induced behavioral plasticity initiated by somatodendritic dopamine release. Indeed, our previous work in adult mice shows that DCC receptor function in VTA dopamine neurons is necessary for the development of AMPH-induced behavioral sensitization^48^.

### DCC receptor upregulation in adolescence prevents AMPH-induced changes in adult PFC dopamine release in adulthood

The fiber photometry recordings revealed subtle changes in baseline dopamine activity in the PFC following adolescent AMPH exposure in those animals transfected with LacZ sgRNA and *Dcc* sgRNA (Fig. 5C-F), but these changes did not reached statistical significance.

Unlike in our previous fiber photometry experiment, we could not reliably detect a peak in response to the saline injection. In fact, in the animals that were transfected with the *Dcc* sgRNA, only 3 out of 10 mice (30%) showed a saline-induced peak. This contrasts with the *LacZ* sgRNA group, in which 9 out of 11 animals showed the stress-induced peak (82%). Although this difference is intriguing, we don’t know why this is the case.

Upon methylphenidate challenge, mice infected with *LacZ* sgRNA and exposed to AMPH in adolescence showed exaggerated dopamine responsiveness, compared to mice pre-exposed to saline (Fig. 5G). Overexpression of DCC receptors in adolescence completely abolishes this effect, indicating that AMPH-induced downregulation of DCC receptors in adolescence drives dopamine hyper-responsiveness in adulthood, most likely due to the presence of ectopic mesolimbic dopamine axons expressing high levels of DAT+. This female-like protective effect, coupled with the absence of AMPH-induced place preference, highlights DCC overexpression as a promising therapeutic target to mitigate short- and long-term consequences of exposure to recreational-like doses of AMPH in adolescence. Immunohistochemistry confirmed robust sgRNA expression within the VTA, co-expressed in TH+ neurons (Fig. 5H).

## Discussion

Misuse of stimulants drugs – both illicit and prescribed – is a growing concern^49^ due to its association with heightened psychiatric risk, including substance use disorders^50–53^, increase suicidal ideation^53^, and depression ^51^. The underlying mechanisms and the causes of the observed differences in vulnerability between men and women during adolescence remain unknown. In male mice, AMPH in early adolescence reduces the expression of the guidance cue receptor DCC in mesolimbic dopamine axons, disrupting their ongoing targeting in the NAc and inducing their ectopic growth to the PFC^11^. Females, however, are protected against these effects. In this study, we assessed the phenotype and function of dopamine axons in the adult PFC of male and female mice exposed to AMPH in adolescence. We found that rewarding, recreational-like doses of AMPH in adolescence lead to exaggerated density of DAT+ fibres in the PFC of adult males, indicating that misrouted mesolimbic dopamine axons retain molecular and functional properties of their intended target ^20–22,54,55^. Indeed, our findings show that adult PFC dopamine signaling in male mice retain mesolimbic-like characteristics, including larger baseline dopamine transients, as well as exaggerated release in response to acute stress and stimulants. Restoring DCC receptor levels in adolescent males via CRISPRa-targeted gene therapy prevents disruptions in dopamine development. In contrast, the same rewarding AMPH doses in females do not alter PFC dopamine phenotype in adulthood, highlighting the male-specific vulnerability to dopamine development disruption in early adolescence by drugs of abuse.

The increase in DAT expression in PFC dopamine terminals likely mediates the alterations in dopamine signaling observed at baseline and in response to acute challenges in male mice. The DAT protein has been shown to regulate the timing, strength, and overall function of dopamine signaling in the striatum and cultured neurons^56^, including the reuptake and repackaging of dopamine after being released^57^. Increased DAT levels in the striatum have been shown to enhance dopamine recycling and reserve pools, reducing the use of newly synthesized dopamine for release ^58^. The larger PFC dopamine transients observed at baseline in AMPH-pretreated male mice compared to saline controls reflect heightened dopamine reuptake, reserve pools, and recycling due to greater PFC DAT+ expression. The fact that the dopamine transients are more sporadic may result from increased dopamine autoreceptor activation, which would decrease neuronal excitability and reduce the probability of dopamine release under baseline conditions ^59^.

Dopamine function in the PFC is highly sensitive to stress^60,61^; we were able to measure group differences in stress-induced changes in dopamine transients via an acute i.p. injection of saline. The steeper and faster dopamine release observed in response to saline in AMPH-pretreated male mice, compared to saline counterparts, is consistent with these mice having greater PFC DAT expression and, therefore, increased dopamine reserve pools. The fact that we did not capture differences between groups in stress-induced dopamine clearance is likely due to the slow signal decay of the GrabDA2h sensor, which cannot detect differences in fast dopamine clearance events^40,41^.

The exaggerated increase in PFC dopamine signal following the methylphenidate challenge in males treated with AMPH in adolescence is also consistent with the presence of DAT+ ectopic mesolimbic dopamine axons in this region and indicates increased potency of methylphenidate. Heightened dopamine function has been observed in the PFC of adult rats overexpressing DAT(DAT-tg)^62^ and, in response to methylphenidate, in the NAc of adult transgenic mice overexpressing DAT(DAT-tg)^63^. Elevated DAT expression in the NAc of rats with a history of methylphenidate self-administration also leads to enhanced drug-induced dopamine release^42,63^. Methylphenidate not only binds to the DAT, but also alters the nerve terminal machinery responsible for dopamine storage and release. It binds to the vesicular monoamine transporter 2 (VMAT2) – a protein that packages extracellular and intracellular dopamine into cytoplasmic and membrane-bound vesicles for subsequent release – thereby increasing the amount of dopamine^64,65^. The increased density of DAT in the PFC of males exposed to AMPH in adolescence likely amplifies this effect by clearing dopamine from the synapse and generating a concentration gradient that favors further dopamine release. The changes in PFC dopamine function due to early adolescent AMPH administration can then be seen as a risk factor leading to drug abuse, not only by decreasing behavioral inhibition^15,19^, but also by increasing the potency of methylphenidate, in addition to potentially other drugs of abuse that increase dopamine release^66^.

In contrast to males, female mice exposed to AMPH in adolescence did not exhibit elevated DAT levels in the PFC or altered dopamine at baseline, or in response to saline or methylphenidate. This sexual dimorphism indicates that PFC dopamine dynamics in males results from ectopic mesolimbic dopamine innervation. Unlike males, females are resilient to the disruptive epigenetic effects of early adolescent AMPH^15^ and do not show *Dcc* mRNA reduction. By mid-adolescence, females show epigenetic changes and *Dcc* mRNA reduction but also exhibit compensatory increases in the DCC receptor ligand, Netrin-1, maintaining normative mesolimbic dopamine axon targeting^15^.

Expression of DAT in adult mice exposed to saline in adolescence is significantly larger in females compared to males – a sex difference that has been reported in the ventral and dorsal striatum of mice^67^ and humans^68,69^. Why adult females have higher PFC DAT levels than males at baseline, is a question that needs to be addressed, but this likely involves sexual dimorphisms in the organization of mesocorticolimbic dopamine circuitry^70^. In contrast with adult male rats, females have a larger proportion of VTA dopamine neurons projecting to the PFC ^71^, as well as more dopamine neurons and greater VTA volume^72^. Interestingly, adult males and ovariectomized adult female rats have lower levels of DAT expression in the VTA and striatum, compared to intact females^73–76^. Despite the overall increase in PFC DAT in female mice, we did not observe sex differences in dopamine dynamics. One reason could be that there are sex differences in mechanisms regulating dopamine release in the PFC. While blockade of NMDA receptors in adult male rats increases extracellular dopamine levels in the PFC^77^, this manipulation reduces dopamine release in females^78^.

Both male and female mice exhibited a robust and long-lasting AMPH conditioned place preference, indicating similar rewarding and positive drug associations^79^. Additionally, both sexes showed an incubation effect, suggesting that the neuroanatomical changes underlying this process are similarly affected in males and females. Resilience in females to AMPH-induced disruption of dopamine development does not imply vulnerability in other domains contributing to addiction. Indeed, while the initial positive experience with the drug may be similar between sexes, the long-term consequences and vulnerability to addiction may diverge. Preclinical models of addiction show that, in general, female rats acquire the self-administration of drugs and escalate their drug taking with extended access more rapidly, show more motivational withdrawal, and show greater reinstatement^80^. Females also show more motivation to work for AMPH^81^. In humans, a similar pattern is observed. While men are more likely than women to use almost all types of illicit drugs^82^, women tend to escalate more rapidly than men do, and once addicted, it is more difficult for females to cease drug use.

Increasing *Dcc* gene expression in the VTA of male mice during adolescence, using CRISPRa, prevented the exaggerated dopamine response to methylphenidate in adulthood. This supports the idea that the mesolimbic-like dopamine phenotype in the PFC of these mice is caused by the rerouting of NAc dopamine axons to the PFC. Restoring DCC levels also prevented the expression of conditional place preference, suggesting changes in reward sensitivity to AMPH. This effect may be mediated by DCC receptor modulation of behavioral plasticity resulting from somatodendritic dopamine release. This idea is supported by our previous work demonstrating that DCC receptor function in the VTA is required for the development of AMPH-induced behavioral sensitization in adult mice^48^. DCC has also been shown to play a critical function in adult brain plasticity, via the orchestration of neuronal circuitry reorganization^83^.

In this study, we demonstrated that AMPH exposure during adolescence in mice leads to male-specific alterations in dopamine signaling in the adult PFC. This effect is due to the mistargeting of NAc dopamine axons to the PFC. These axons adopt molecular and functional properties of mesolimbic-projecting dopamine neurons, including increased DAT expression and exaggerated responses to acute MPH. Dopamine neurotransmission in the PFC plays a crucial role in higher-order cognition^84–88^. The alterations in PFC dopamine signaling induced by AMPH in adolescence may underlie the risk for cognitive deficits observed in disorders of poor impulse control, including drug addiction^89–93^. Our results indicate that DCC receptor upregulation, via pharmacological or non-invasive innervation, may be a promising target for therapeutic intervention^94–96^.

## Supporting information

Stats tables

## Acknowledgments

This work was supported by the National Institute on Drug Abuse at the National Institutes of Health (R01DA037911 to C.F), the Canadian Institutes of Health Research (PJT 190045; FRN:17030, 170130 to C.F.), and the Natural Sciences and Engineering Research Council of Canada (RGPIN-2020-04703 to C.F; CGS-M to T.C.).

The present study used the services of the Molecular and Cellular Microscopy Platform in the DHRC. Melina Jaramillo Garcia and Bita Khadivjam helped set up the imaging experiments. We thank Dr. Nicolas Tritsch and Dr. Jorge Quillfeldt for their insightful comments on the manuscript.

## Author Contributions

GH and CF designed the study. GH, MS, and SG performed behavioral experiments. GH, JZ, DM, and TC performed fibre photometry experiments. JZ, ZN, TC, AM, IH, and GH performed neuroanatomy experiments. JJD provided CRISPRa virus material and the sgRNAs. GH and CF analyzed the data. GH and CF wrote the manuscript. All authors discussed the results, edited, and approved the manuscripts

